# Abbreviated assessment of psychopathology in patients with suspected seizure disorders

**DOI:** 10.1101/677278

**Authors:** Charles B Malpas, Albert D Wang, Michelle Leong, Benjamin Johnstone, Genevieve Rayner, Tomas Kalincik, Patrick Kwan, Terence J O’Brien, Dennis Velakoulis

## Abstract

**Purpose:** Psychopathology is common in patients undergoing investigation for seizure-related disorders. Psychometric examination using self-report instruments, such as the SCL-90-R, can assist diagnosis. The SCL-90-R, however, is a lengthy instrument and might not be tolerated by all patients. We assessed several abbreviated forms of the SCL-90-R in patients undergoing video encephalographic monitoring (VEM).

**Method:** 687 patients completed the SCL-90-R and scores were computed for the full SCL-90-R and five abbreviated forms. Correlations and mean differences were computed between different forms. Classification accuracy was assessed via receiver operating characteristic (ROC) curves, and measurements models were examined using confirmatory factor analysis (CFA).

**Results:** All abbreviated forms were strongly correlated with the SCL-90-R for general psychopathology (*r* = .93 – .99), depression (*r* = .89 – .95), anxiety (*r* = .97 – .98), psychosis (*r* = .95 – .99), and obsessive-compulsive symptoms (*r* = .97). Classification performance was similar across forms for depression and anxiety, with high negative predictive values (.90 – .94) and lower positive predictive values (.34 – .38). Classification performance for psychotic and obsessive-compulsive disorders was poor. Differences were observed between the full SCL-90-R and its abbreviated forms across most domains (*d* = 0.00 – 0.65). The published measurement model was most strongly validated for the SCL-27, SCL-14, and the SCL-K-9.

**Conclusions:** These five SCL-90-R abbreviated forms show high convergent validity with the full version. In patients undergoing investigation for seizure-related disorders, the BSI or BSI-18 are most appropriate where screening for both depression and anxiety is required. The SCL-K-9 is appropriate when only a single measure of global psychological distress is required. None of the instruments were able to detect psychotic or obsessive-compulsive symptoms with great accuracy. Caution should be exercised when making direct comparisons across the different forms.

## 1. Introduction

Psychiatric comorbidity is common in patients with epilepsy and related disorders. It is estimated that up to 30% of patients with epilepsy meet criteria for a major psychiatric disorder, and rates of depressive and anxiety related disorders in particular are high compared to the general population (Karouni et al., 2010; Rai et al., 2012; Tellez-Zenteno, Patten, Jetté, Williams, & Wiebe, 2007; Weatherburn, Heath, Mercer, & Guthrie, 2017). The rate of comorbid psychopathology is also high in patients with psychogenic non-epileptic seizures. Between 53 and 100 percent of patients with PNES are estimated to meet criteria for an additional major psychiatric disorder, with notable elevations in the rates of post-traumatic, personality, anxiety, and depressive disorders relative to patients with epilepsy (Diprose, Sundram, & Menkes, 2016). The rates of psychopathology in patients undergoing video encephalographic monitoring (VEM) are likely higher than the general epilepsy population, with up to 50% of patients meeting criteria for a comorbid major psychiatric disorder (Blumer, Montouris, & Hermann, 1995). VEM patients, in particular, are highly likely to have disruption of key cognitive networks underpinning depressive symptomatology (Rayner, 2017). Taken together, there is a clear and present need to assess psychiatric comorbidities in patients undergoing investigation for epilepsy.

While full neuropsychiatric assessment during VEM admission is the gold-standard for psychiatric diagnosis, this process is time consuming, costly, and might not be available in all centres (Agrawal, Turco, Goswami, Faulkner, & Singh, 2015). The use of valid, reliable, and comprehensive psychometric instruments provides a practical approach to screen for psychiatric symptomatology, thus facilitating efficient use of formal neuropsychiatric assessment (de Oliveira et al., 2014; Jones et al., 2005). The Symptom Checklist 90 Revised (SCL-90-R) is a comprehensive self-report instrument of psychiatric symptomatology, consisting of 90 questions that takes approximately 20 minutes to complete (Derogatis, 1992). Unlike many brief screening instruments, the SCL-90-R covers nine psychiatric domains, making it useful for detecting a broad range of psychopathologies. A large body of evidence has investigated the reliability and validity of the SCL-90-R across a number of settings (Brophy, Norvell, & Kiluk, 1988; Derogatis & Savitz; Rauter, Leonard, & Swett, 1996; Schmitz et al., 2000; Wiebe, Rose, Derry, & McLachlan, 1997).

The length of the SCL-90-R, however, might make it impractical in some clinical contexts. To address this, several abbreviated forms of the SCL-90-R have been developed. These include the SCL-27 (27 items; Hardt & Gerbershagen, 2001), SCL-17 (14 items; Harfst et al., 2002), SCL-K-9 (9 items; Klaghofer & Brähler, 2001), BSI (53 items; Derogatis, 1993), and BSI-18 (18-items; Derogatis, 2000). Several studies have shown that these abbreviated forms show excellent correspondence with the long form SCL-90-R in general psychiatric populations (Müller, Postert, Beyer, Furniss, & Achtergarde, 2010; Prinz et al., 2013; Sereda & Dembitskyi, 2016). The BSI, in particular, has been shown to be an appropriate substitute for the SCL-90-R when time limitation is a factor. No studies, however, have directly compared the SCL-90-R to any of the abbreviated forms in a population of patients undergoing investigation for presumed seizure-related disorders. There is a paucity of research comparing these different forms amongst English-speaking patients. As such, there is no strong evidence to inform decisions regarding the use of these abbreviated instruments in VEM settings.

An understanding of the relationship between the SCL-90-R and its short forms is also necessary to interpret the existing epilepsy literature that uses these instruments. For example, the BSI has been used to study the prevalence of psychiatric morbidity in general patients with epilepsy, as well as those with concomitant epilepsy and intellectual disability (Endermann, 2005; Swinkels, Kuyk, Van Dyck, & Spinhoven, 2005). The BSI-18 has been used to investigate similarities between psychogenic seizure disorders and psychogenic movement disorders, and also to determine the overlap between dissociation and other psychiatric symptoms in patients experiencing psychogenic seizures (M. L. Cohen, Testa, Pritchard, Zhu, & Hopp, 2014; Hopp, Anderson, Krumholz, Gruber-Baldini, & Shulman, 2012). An implicit assumption underpinning the use of these short forms is that they correspond to the established psychometric properties of the full SCL-90-R in epilepsy populations.

The overall aim of this study was to directly compare the SCL-90-R to its various abbreviated forms in a large sample of patients undergoing VEM. The primary focus was to examine correlations and mean differences across the various instruments. The secondary focus was on the ability for these instruments to detect symptoms of depression, anxiety, psychosis, obsessive-compulsive disorder, and global psychopathology. Following Prinz and colleagues (2013), it was hypothesised that all abbreviated forms would show strong correspondence with the long form, but that some versions would show better classification performance when benchmarked against gold-standard neuropsychiatric diagnosis. Using confirmatory factor analysis, we also expected to find evidence for the measurement models proposed by the test developers for the various forms of the SCL-90-R.

## 2. Material and Methods

### 2.1 Participants

Participants were patients admitted for prolonged VEM at the Royal Melbourne Hospital, Melbourne, Australia between 2002 and 2017. Patients were eligible if they: (a) had undergone VEM and formally been discussed during the consensus meeting, (b) had undergone formal neuropsychiatric evaluation during admission, (c) completed the SCL-90-R during admission, (d) were aged 18 years or over at the time of admission, and (e) had sufficient English language proficiency to complete the SCL-90-R and other psychometric instruments. This study was approved by the Melbourne Health Ethics committee (HREC# QA2012044).

### 2.2 Epilepsy diagnosis

VEM consensus diagnosis was made by a multidisciplinary team meeting of neurologists, neuropsychiatrists, neuropsychologists, neuroradiologists, and neurophysiology scientists.

The patient’s clinical history, VEM results, neurological examination, neuropsychological profile, neuropsychiatric assessment, and neuroimaging findings (where applicable) were all considered when formulating a final diagnosis. Epilepsy was defined as the presence of two or more unprovoked seizures or one seizure with enduring risk for seizure occurrence of >60% (Fisher et al., 2014). Psychogenic non-epileptic seizures (PNES) were defined as sudden, involuntary changes in behaviour, sensation and/or motor activity, with or without alterations in conscious state, associated with psychosocial distress and without electrographic correlate (Lesser, 1996). For the purposes of this research, patients were classified into one of five categories: epilepsy, PNES, concomitant epilepsy and PNES, other non-epileptic events, and non-diagnostic.

### 2.3 Psychiatric diagnosis

All patients admitted for VEM underwent formal psychiatric assessment by a specialist neuropsychiatrist. Diagnosis was informed by the Diagnostic and Statistical Manual of Mental Disorders 4^th^ edition (DSM-IV-TR; American Psychiatric Association, 2000) or 5^th^ edition (DSM-5; American Psychiatric Association, 2013). Patients were categorised as currently having: any major psychiatric disorder (axis 1 or equivalent), any depressive disorder, any anxiety disorder, any psychotic disorder, or obsessive-compulsive disorder. While other diagnoses were made, the relatively frequency was low and they did not directly map onto the psychopathology domains measured by the SCL-90-R (e.g., eating disorders, attention-deficit hyperactivity disorder. SCL-90-R scores were not known to the clinician during diagnostic formulation. These categories were chosen because they represent the major psychiatric diagnoses observed in this population. Depression was chosen, in particular, because of its association with suicidal ideation and behaviour, and the need to detect these symptoms and provide early intervention and assessment.

### 2.4 Psychometric instruments

Patients were administered the long form Symptom Checklist 90 Revised (SCL-90-R; Derogatis, 1992), which is a 90-item self-report inventory of psychological symptoms. The SCL-90-R measures symptomatology along nine domains: somatisation, obsessive compulsive symptoms, interpersonal sensitivity, depression, anxiety, hostility, phobic anxiety, paranoid ideation, and psychoticism. Participants respond to the stem ‘for the past week, how much were you bothered by:’ on a 5-point Likert scale from 0 (*Not at all*) to 4 (*Extremely*). Each domain is scored by summing the items and dividing by the number of non-missing responses. A global severity index (GSI) is computed by summing all 90 items and dividing by the number of non-missing responses.

Scores for five abbreviated forms of the SCL-90-R were computed, each using a subset of items from the full SCL-R-90. These included the SCL-27 (27 items; Hardt & Gerbershagen, 2001), SCL-17 (14 items; Harfst et al., 2002), SCL-K-9 (9 items; Klaghofer & Brähler, 2001), BSI-53 (53 items; Derogatis, 1993), and BSI-18 (18-items; Derogatis, 2000). As with the long form SCL-90-R, each domain was scored by summing the items and dividing by the number of non-missing responses. A GSI was computed for each abbreviated form.

A subset of patients also completed the Hospital Anxiety and Depression Scale (HADS; Zigmond & Snaith, 1983), which is a 14-item screening instrument for depression and anxiety. Patients rate each item on a 4-point Likert scale ranging from 0 to 3. The scores are summed across 7 items each for the depression and anxiety domains to form subscale scores.

### 2.5 Statistical analyses

All analyses were performed in R (R Core Team, 2019). Cohen’s (1988) *d* was computed to examine the difference between full SCL-90-R and abbreviated form test scores, and to compare scores between diagnostic groups. Pearson’s (1895) product moment correlation coefficient (*r*) was computed to examine the correlation between full SCL-90-R and abbreviated form test scores. Classification performance was examined by computing receiver operating characteristic (ROC) curves using the *pROC* package (Robin et al., 2011). The area under the curve (AUC) was computed. An optimal cut-off was determined using Youden’s (1950) *J* statistic, and the sensitivity, specificity, negative predictive values (NPV), and positive predictive value (PPV) computed. Statistical inference was performed via 95% confidence intervals.

Confirmatory factor analysis (CFA) was performed using the *lavaan* package (Rosseel, 2012). Oblique models were specified as per the structure proposed by the test authors. Cross-loadings and residual covariances were fixed to zero. Robust estimation was performed using full-information maximum likelihood (ML) with Huber-White standard errors and scaled test statistic that asymptotically approach the Yuan-Bentler statistic. Reliability was computed for the latent variables using the omega coefficient (formula 21; Green & Yang, 2009). Model fit was determined using the following robust indices: *χ*^2^ (Jöreskog, 1969), root mean squared error of approximation (RMSEA; Steiger & Lind, 1980), standardised root mean residual (SRMR; Bentler, 1995), Tucker-Lewis index (TLI; Tucker & Lewis, 1973) and the comparative fit index (CFI; Bentler, 1990). Acceptable fit was considered to be: RMSEA < .06, SRMR < .08, TLI > .95, and CFI > .95 (West, Taylor, & Wu, 2012).

## 3. Results

### 3.1 Sample characteristics

Of the original 687 patients with available demographic, clinical, and psychometric data, 611 (89%) had ≤ 20% missing data on the SCL-90-R and were included in the final sample. The majority of patients had no missing SCL-90-R items (*n* = 545, 89%). 36 patients had one missing item (6%), 16 had two missing items (3%), and seven participants had between three and 12 missing items (2%). There was no obvious pattern to the missing items. The majority of participants were female (*n* = 387, 63%) and the mean age was 39 (*SD* = 15, *Range* = 18 - 87). The most common diagnosis was epilepsy (*n* = 286, 47%), followed by psychogenic non-epileptic seizures (n = 151, 25%), other non-epilepsy events (*n* = 34, 6%), and concomitant epilepsy and psychogenic non-epilepsy seizures (*n* = 33, 5%). A diagnosis could not be reached for 107 patients (18%). Neuropsychiatric assessment revealed that 209 (34%) patients met criteria for a current major psychiatric disorder. Of all patients, 126 (21%) met criteria for a depressive disorder, while 122 (20%) met criteria for an anxiety disorder.

### 3.2 Differences and similarities between long and short forms

*Table 1* shows the relationship between the SCL-90-R scores and the relevant scores on the abbreviated forms. A GSI is shown for all the abbreviated forms, and scores are also shown for the depression, anxiety, psychoticism, paranoia, and obsessive-compulsive domains where the short form has a comparable scale. As shown in *Figure 1*, there was a high concordance between the GSI across difference forms of the test. The relationships were slightly weaker for the depression, anxiety, psychoticism, and obsessive-compulsive domains (*Figure S1-S3*). The correlation between the SCL-90-R scores and the short form scores was very high, ranging between .89 for the depression score of the SCL-27 to .99 for the GSI on the BSI. Nevertheless, differences between mean scores were observed for most scales. The greatest difference was between the SCL-27 depression score and the SCL-90-R depression score, which was a medium to large effect size (*d* = .65) by Cohen’s estimation (1988).

**Figure 1.**
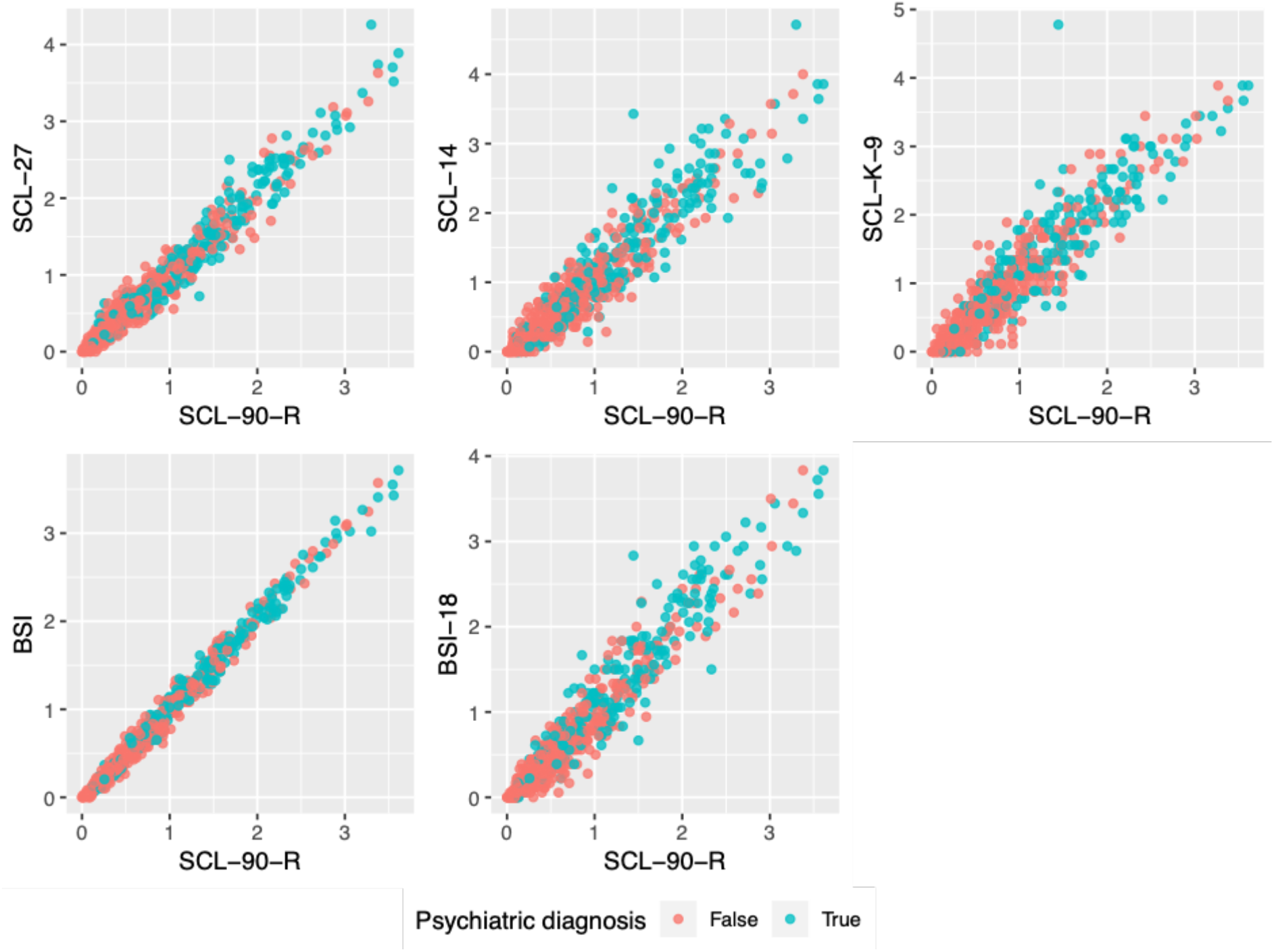
Correlations between the Global Severity Index (GSI) of the long form SCL-90-R and abbreviated forms. As shown, the correlations between the measures was high (*r* = .93 - .99). Patients with any major psychiatric diagnosis are shown in green, while those without any diagnosis are shown in red.

**Figure 2.**
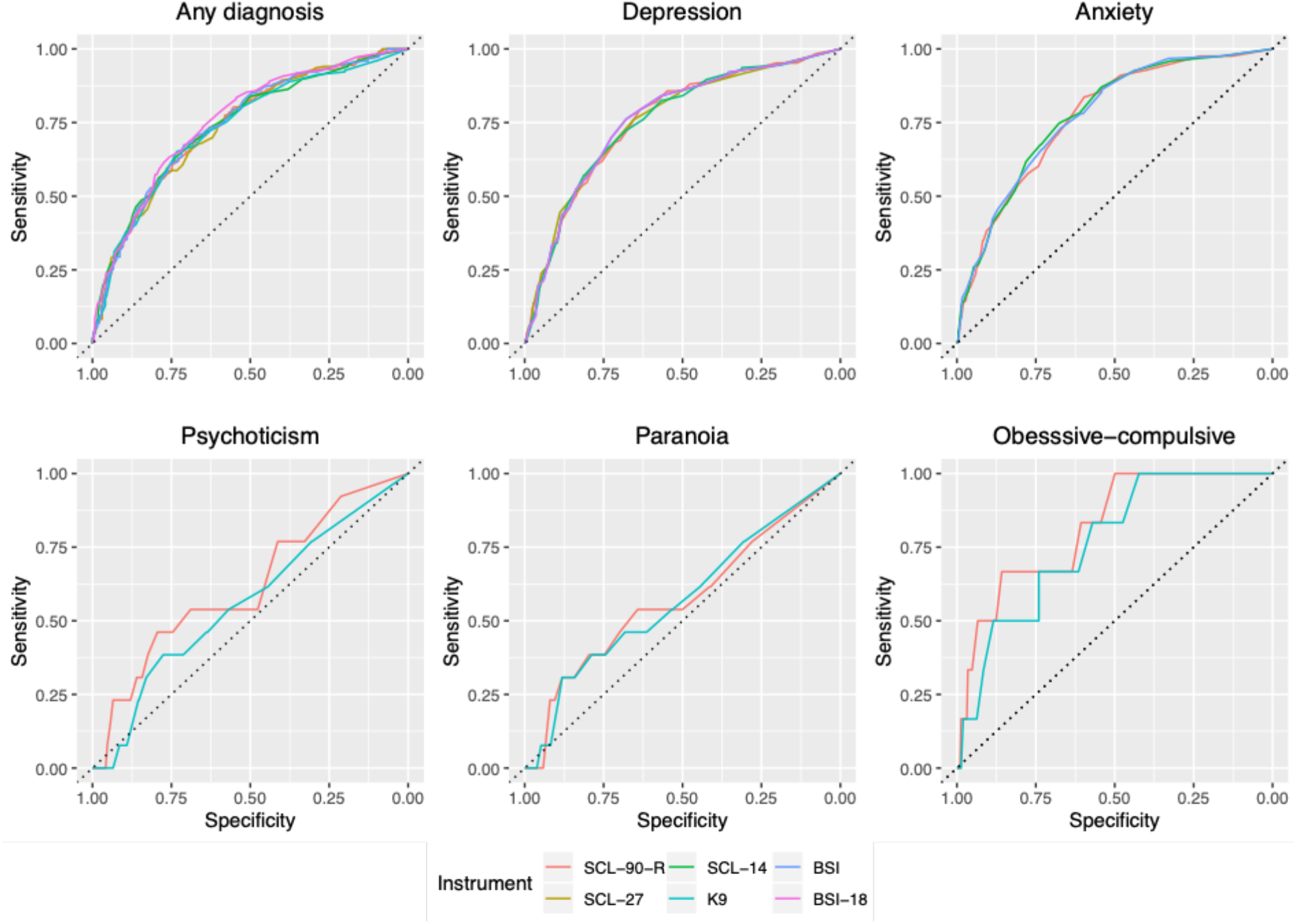
Receiver operating characteristic (ROC) curves for the SCL-90-R and its abbreviated forms. The top row shows comparable classification performance for any diagnosis, depression or anxiety. The bottom row shows the poorer performance for psychosis (using the psychoticism and paranoia scores) and obsessive-compulsive disorder.

**Table 1.** Relationship between SCL-90 and short forms

|  | *M<sub>diff</sub>* [95% CIs] | *d* [95% CIs] | *r* [95% CIs] |
| --- | --- | --- | --- |
| *GSI* |  |  |  |
| SCL-27 | **0.02 [0.01, 0.03]** | **0.12 [0.04, 0.20]** | **.98 [.97, .98]** |
| SCL-14 | **0.04 [0.02, 0.06]** | **0.14 [0.06, 0.22]** | **.94 [.93, .95]** |
| SCL-K-9 | **0.09 [0.07, 0.12]** | **0.27 [0.19, 0.35]** | **.93 [.92, .94]** |
| BSI | **0.01 [0.12, 0.17]** | **0.10 [0.02, 0.18]** | **.99 [.99, .99]** |
| BSI-18 | 0.00 [-0.02, 0.02] | 0.00 [-0.08, 0.08] | **.96 [.95, .96]** |
| *Depression* |  |  |  |
| SCL-27 | **0.29 [0.25, 0.33]** | **0.65 [0.56, 0.73]** | **.89 [.87, .90]** |
| SCL-14 | **0.07 [0.04, 0.09]** | **0.21 [0.13, 0.29]** | **.95 [.94, .96]** |
| BSI | **0.15 [0.12, 0.17]** | **0.47 [0.39, 0.55]** | **.95 [.94, .96]** |
| BSI-18 | **0.15 [0.12, 0.17]** | **0.47 [0.39, 0.55]** | **.95 [.94, .96]** |
| *Anxiety* |  |  |  |
| BSI | **0.07 [0.05, 0.09]** | **0.32 [0.24, 0.40]** | **.97 [.97, .98]** |
| BSI-18 | **0.05 [0.04, 0.07]** | **0.27 [0.19, 0.35]** | **.98 [.97, .98]** |
| *Psychoticism<sup>#</sup>* |  |  |  |
| BSI | **0.07 [0.05, 0.09]** | **0.09 [0.06, 0.11]** | **.95 [.94, .96]** |
| *Paranoia<sup>#</sup>* |  |  |  |
| BSI | **0.01 [0.01, 0.03]** | 0.02 [0.00, 0.03] | **.99 [.98, .99]** |
| *Obsessive-compulsive* |  |  |  |
| BSI | **0.18 [0.16, 0.20]** | **0.17 [0.15, 0.19]** | **.97 [.96, .97]** |

**Note:** *M<sub>diff</sub>* is the absolute mean difference between two groups on the relevant measure. *d* = is Cohen's *d*, a standardised measure of effect size. *r* = Pearson's product moment correlation coefficient between the SCL-90-R subscale and the equivalent subscale from the abbreviated form. 95% confidence intervals are shown. Bold indicates confidence intervals that do not capture 0. <sup>#</sup> = Diagnostic accuracy of both the PAR (paranoia) and PSY (psychoticism) scales were determined using the clinical diagnosis of any current psychotic disorder.

### 3.3 Convergent validity

The majority of patients (*n* = 593, 97%) completed the HADS, which was used to examine the convergent validity of the SCL-90-R and abbreviated forms. The HADS depression score was highly correlated with the depression scores from the SCL-90-R (*r* = .70 [95%CI = .66, .74]), SCL-27 (*r* = .63 [95%CI = .58, .67]), SCL-14 (*r* = .68 [95%CI = .63, .72]), BSI (*r* = .68 [95%CI = .63, .72]), and the BSI-18 (*r* = .68 [95%CI = .63, .72]). Similarly, the HADS anxiety score was highly correlated with the anxiety scores on the SCL-90-R (*r* = .75 [95%CI = .72, .79]), BSI (*r* = .76 [95%CI = .72, .79]), and the BSI-18 (*r* = .76 [95%CI = .72, .79]).

### 3.4 Criterion validity

The mean differences between psychiatric diagnostic groups was used to evaluate the criterion validity of the SCL-90-R and its short forms. *Table 2* shows the mean differences on GSI scores between patients with any major psychiatric diagnosis and those without. Also shown are mean differences on depression scores between patients with and without a depressive diagnosis, and mean differences on anxiety scores between patients with and without an anxiety diagnosis. Mean differences are shown on both the psychoticism and paranoia scales for patients with and without a current psychotic disorder diagnosis. Finally, the mean differences are shown on the obsessive-compulsive scale for patients with and without a diagnosis of obsessive-compulsive disorder. As shown, all instruments showed large (*d* > .80) differences between groups for depression, anxiety, and obsessive-compulsive disorder. There was no strong evidence for mean differences in the paranoia and psychoticism scales.

**Table 2.** Differences between diagnostic groups

| | $M_{diff}$ [95% CIs] | $d$ [95% CIs] |
| --- | --- | --- |
| *Any diagnosis* |  |  |
| SCL-90-R | **0.61 [0.49, 0.73]** | **0.87 [0.66, 1.02]** |
| SCL-27 | **0.64 [0.51, 0.77]** | **0.86 [0.64, 1.00]** |
| SCL-14 | **0.73 [0.59, 0.87]** | **0.89 [0.67, 1.03]** |
| SCL-K-9 | **0.72 [0.57, 0.86]** | **0.85 [0.64, 1.00]** |
| BSI | **0.63 [0.51, 0.76]** | **0.87 [0.66, 1.02]** |
| BSI-18 | **0.71 [0.58, 0.85]** | **0.94 [0.72, 1.08]** |
| *Depression* |  |  |
| SCL-90-R | **0.85 [0.66, 1.03]** | **0.96 [0.70, 1.13]** |
| SCL-27 | **0.88 [0.68, 1.09]** | **0.91 [0.63, 1.06]** |
| SCL-14 | **0.92 [0.72, 1.12]** | **0.94 [0.69, 1.13]** |
| BSI | **0.93 [0.73, 1.13]** | **0.95 [0.69, 1.13]** |
| BSI-18 | **0.93 [0.73, 1.13]** | **0.95 [0.69, 1.13]** |
| *Anxiety* |  |  |
| SCL-90-R | **0.79 [0.61, 0.97]** | **0.94 [0.66, 1.10]** |
| BSI | **0.88 [0.69, 1.07]** | **0.98 [0.70, 1.14]** |
| BSI-18 | **0.85 [0.67, 1.04]** | **0.98 [0.70, 1.15]** |
| *Psychoticism<sup>#</sup>* |  |  |
| SCL-90-R | 0.26 [-0.11, 0.63] | 0.39 [-0.16, 0.94] |
| BSI | 0.11 [-0.32, 0.54] | 0.13 [-0.41, 0.69] |
| *Paranoia<sup>#</sup>* |  |  |
| SCL-90-R | 0.24 [-0.21, 0.68] | 0.29 [-0.26, 0.84] |
| BSI | 0.22 [-0.23, 0.68] | 0.27 [-0.28, 0.82] |
| *Obsessive-compulsive* |  |  |
| SCL-90-R | **1.19 [0.44, 1.95]** | **1.28 [0.47, 2.09]** |
| BSI | **1.01 [0.17, 1.85]** | **0.97 [0.16, 1.78]** |

**Note:** $M_{diff}$ is the mean difference between two groups on the relevant measure. $d$ is Cohen's $d$ , a standardised measure of effect size. 95% confidence intervals are shown. Bold indicates confidence intervals that do not capture 0. <sup>#</sup> = Diagnostic accuracy of both the PAR (paranoia) and PSY (psychoticism) scales were determined using the clinical diagnosis of any current psychotic disorder.

### 3.5 Classification performance

Receiver operating characteristic (ROC) curves were computed to evaluate the ability of the SCL-90-R and its short forms to classify patients according to psychiatric diagnosis. As shown in *Table 3*, all instruments demonstrated statistically significant areas under the curve (AUC) for depression, anxiety, and obsessive-compulsive disorder, suggesting that each was sensitive to the relative diagnostic group. The AUCs were not significant for psychosis, either in terms of the paranoia or psychoticism scales. The sensitivity, specificity, negative predictive value (NPV), and positive predictive value (PPV) are also shown based on an optimal cut-off score derived using Youden’s (1950) *J* statistic. As shown, the sensitivities and specificities were generally low (< .80) across measures. The only exception was the anxiety score from the SCL-90-R which exhibited a sensitivity of .84 for detecting patients with an anxiety disorder. The PPVs were relatively low across measures (.34 - .58), indicated a relatively low probability that a patient screening positively would subsequently meet diagnostic criteria. The NPVs, however, were much higher (.80 - .94) suggesting that a negative screen was significantly predictive of a patient not meeting criteria for a psychiatric diagnosis. As shown in *Figure 1*, the ROC curves were comparable across instruments, suggesting little difference in their ability to classify patients in terms of psychiatric diagnosis.

**Table 3.** Classification performance

|  | *AUC [95%*<br>*CI]* | *Cut-off* | *Sensitivity* | *Specificity* | *NPV* | *PPV* |
| --- | --- | --- | --- | --- | --- | --- |
| *Any diagnosis* |  |  |  |  |  |  |
| SCL-90-R | **.75 [.71, .79]** | 0.81 | .72 | .67 | .82 | .53 |
| SCL-27 | **.74 [.70, .78]** | 0.83 | .67 | .69 | .80 | .53 |
| SCL-14 | **.75 [.71, .79]** | 0.94 | .65 | .74 | .80 | .56 |
| SCL-K-9 | **.73 [.70, .78]** | 1.06 | .67 | .73 | .81 | .57 |
| BSI | **.75 [.71, .79]** | 0.86 | .69 | .71 | .82 | .56 |
| BSI-18 | **.76 [.72, .80]** | 0.92 | .65 | .76 | .81 | .58 |
| *Depression* |  |  |  |  |  |  |
| SCL-90-R | **.76 [.71, .81]** | 1.04 | .79 | .64 | .92 | .36 |
| SCL-27 | **.76 [.71, .81]** | 0.71 | .76 | .75 | .91 | .36 |
| SCL-14 | **.76 [.71, .81]** | 1.08 | .72 | .68 | .90 | .37 |
| BSI | **.76 [.71, .81]** | 0.92 | .76 | .68 | .92 | .38 |
| BSI-18 | **.76 [.71, .81]** | 0.92 | .76 | .68 | .92 | .38 |
| HADS-D | **.77 [.72, .82]** | 7.50 | .64 | .82 | .90 | .48 |
| *Anxiety* |  |  |  |  |  |  |
| SCL-90-R | **.77 [.72, .81]** | 0.68 | .84 | .60 | .94 | .34 |
| BSI | **.77 [.73, .81]** | 0.92 | .75 | .67 | .91 | .36 |
| BSI-18 | **.77 [.71, .81]** | 0.92 | .75 | .67 | .91 | .36 |
| HADS-A | **.71 [.65, .76]** | 9.50 | .64 | .73 | .89 | .38 |
| *Psychosis<sup>#</sup>* |  |  |  |  |  |  |
| SCL-90-R | .62 [.46, .78] | 0.95 | .46 | .79 | .99 | .05 |
| BSI | .56 [.40, .73] | 1.10 | .38 | .78 | .98 | .04 |
| *Paranoia<sup>#</sup>* |  |  |  |  |  |  |
| SCL-90-R | .57 [.40, .75] | 1.75 | .31 | .88 | .98 | .05 |
| BSI | .57 [.40, .74] | 1.70 | .31 | .88 | .98 | .05 |
| *Obsessive-compulsive* |  |  |  |  |  |  |
| SCL-90-R | **.81 [.66, .98]** | 2.45 | .67 | .86 | 1.00 | .04 |
| BSI | **.77 [.59, .94]** | 1.08 | 1.00 | .42 | 1.00 | .02 |

**Note:** HADS-D = Hospital anxiety and depression scale (HADS) depression score. HADS-A = HADS anxiety score. AUC = area under the curve with 95% confidence intervals computed using DeLong's method (1988) with the approach described by Sun and Xu (2014). Bold indicates AUC intervals that do not capture .5. <sup>#</sup> = Diagnostic accuracy of both the PAR (paranoia) and PSY (psychoticism) scales were determined using the clinical diagnosis of any current psychotic disorder.

The HADS and all SCL-90-R forms performed equally well in terms of diagnostic classification of depression and anxiety. As shown in *Table 3*, the AUCs were comparable for the depression diagnoses, and the SCL-90-R and its abbreviated forms slightly outperformed the HADS for anxiety diagnoses. Direct comparison of the AUCs, however, did not reach statistical significance.

### 3.5 Measurement structure

Confirmatory factor analysis (CFA) was performed to evaluate the measurement structure for the SCL-90-R and the various short forms. As shown in *Table S1*, the various forms of the instrument varied in their fit to the data. The SCL-27, SCL-14, and K9 provided good fit across all indices. The BSI and BSI-18 exhibited marginal fit. While the RMSEA and SRMR were acceptable, the CFI and TLI were slightly below the acceptable range. The SCL-90-R exhibited the worst to the data. Again, while the RMSEA and SRMR were acceptable, the CFI and TLI indicated poor fit to the data. Reliability coefficients for each scale are included in *Table S2*. Reliabilities were high for the global severity indices (GSI) across all instruments (*ω* = .89 - .97). Domain scores showed adequate reliability for the SCL-27 (*ω* = .80 - .85), SCL-14 (*ω* = .75 - .89), BSI (*ω* = .79 - .91), and BSI-18 (*ω* = .81 - .91).

## 4. Discussion

In this large study of patients undergoing investigation of presumed seizure disorders, we directly compared the SCL-90-R to five commonly used abbreviated form measures. Three key findings emerged. First, very high correlations were observed between the short forms and long form in the domains of general psychopathology, depression, anxiety, psychosis, and obsessive-compulsive symptoms. This strongly suggests that the various short forms measure these same constructs as the full SCL-90-R. Second, we confirmed that all forms of the SCL-90-R produced equivalent effect sizes when comparing cases and controls across general psychopathology, depression, anxiety, psychosis, and obsessive-compulsive domains. This suggests that the different short-forms are equally appropriate for detecting differences is psychopathology at the group level in terms of depression, anxiety, and obsessive-compulsive symptoms. Third, the various abbreviated forms showed similar classification performance when compared to gold-standard neuropsychiatric diagnosis of depression, anxiety, and obsessive-compulsive disorder. None of the instruments used in the study were sensitive to psychosis, as measured by either the psychoticism or paranoia psychopathology domains. While group differences were observed for obsessive-compulsive disorder, no instruments demonstrated useful negative or positive predictive values. The utility of the SCL-90-R and its short-forms appears limited for detecting psychotic or obsessive-compulsive symptomatology in patients undergoing VEM.

Taken together, however, these findings suggest that the various abbreviated forms of the SCL-90-R can be substituted for the full version in many contexts. For example, in situations where a continuous outcome measure is required for a clinical trial, any one of the short forms would be appropriate. The determining factor is the desired coverage of the available psychopathology domains. For example, if measurement of all nine domains is required, then the BSI is appropriate because it is the only short form measurement with the same content as the full SCL-90-R. If, however, only depression and anxiety domains are of interest, then either the BSI or BSI-18 short forms are appropriate, as these are the only two short forms that explicitly contain depression and anxiety domains. If a single measure of global psychopathology is required, then any of the short-forms would provide an appropriate substitute. The SCL-K-9, the shortest instrument considered here with only 9 items, was highly correlated with the SCL-90-R’s global severity index. These findings are consistent with previous work in non-epilepsy populations that demonstrated strong inter-instrument correlation between the SCL-90-R and its various short forms (Müller et al., 2010; Prinz et al., 2013; Sereda & Dembitskyi, 2016).

It is important to note that, while all of these instruments correlated highly with the relevant SCL-90-R scale, there were mean differences in test scores. This suggests that, while short forms can be used in place of the long form SCL-90-R, the scores cannot be treated as exchangeable. It would not be appropriate, for example, to compare scores on the long-form SCL-90-R with scores on the short form. A single instrument must be administered to all participants in order for comparisons to be valid.

The various short forms of the SCL-90-R showed similar classification performance compared to clinical diagnosis. It is important to note, however, that the accuracy of these instruments was not optimal for diagnostic purposes. The overall accuracy of these instruments, as indicated by the ROC analyses, was statistically significant, but limited. The low PPVs indicate relatively low probabilities that a patient classified as a ‘positive’ case actually has the relevant disorder. In contrast, the high NPVs indicate that ‘negative’ cases are highly likely to not have the relevant disorder on formal psychiatric assessment. The practical implication is that patients below the relevant cut-off can reasonably be assigned a lower priority for formal psychiatric assessment (Cut-off scores are shown in *Table 3*). These findings are consistent with previous research investigating the screening accuracy of the SCL-90-R in neurological disorders (Aben, Verhey, Lousberg, Lodder, & Honig, 2002; Leathem & Babbage, 2000; Ruis et al., 2014). It is important to note that none of the instruments used in this study showed great sensitivity or specificity to psychotic or obsessive-compulsive disorders.

We also compared the classification performance of the various SCL-90-R forms to the Hospital Anxiety and Depression Scale (HADS), a commonly used screening test for depression and anxiety (HADS; Zigmond & Snaith, 1983). The HADS is a 14-item instrument, which makes it comparable to many of the SCL-90-R short forms in terms of administration time. Our findings indicated that the classification performance of the SCL-90-R and its short forms was comparable to the HADS for both depression and anxiety diagnoses. In this sense, the HADS might be an optimal choice in clinical or research settings where screening is only required for depression or anxiety. The SCL-90-R and its short forms, however, might be preferable in situations where the measurement of other symptom domains is of interest.

It is important to note that the performance of the SCL-90-R and its short forms in screening for depression and anxiety was comparable to other readily available screening instruments. For example, the Neurological Disorders Depression Inventory for Epilepsy (NDDI-E; Gilliam et al., 2006) is a 6-item screening tool for depression. The NDDI-E performs similarly well to the SCL-90-R and its short forms in detecting depression, showing an area under the curve (AUC) of .77, a positive predictive value of .53, and a negative predictive value of .86 in the initial validation study. As such, the NDDI-E might be preferable for clinical or research settings in which only depressive symptomatology is of interest.

Investigation of latent measurement models across the various instruments produced mixed results. For each instrument, we tested the oblique measurement model proposed by the original authors. Correlations between the latent variables were free to correlate. Cross loadings and residual covariances were fixed to zero. This represented the ‘ideal’ measurement model for each instrument. Only three instruments fitted this model adequately, namely the SCL-27, SCL-14, and SCL-K-9. The remaining instruments came close to fitting this model form but produced relative fit statistics that brought the latent variable into question. These findings are consistent with previous work raising questions as to the most appropriate measurement model for the SCL-90-R (Cyr, McKenna-Foley, & Peacock, 1985; Rauter et al., 1996; Urbán et al., 2014; Vassend & Skrondal, 1999).

The practical implications of these findings depend on the specific application. For screening purposes, the appropriateness of the psychometric instrument depends more on the empirical question of classification accuracy and less on the specific latent variable structure of the instrument. As discussed above, for screening of depression and anxiety, there is little difference between the SCL-90-R and its various short forms in VEM populations. In situations where the accurate measurement of well-defined psychopathological traits is required, the SCL-27, SCL-14, and SCL-K-9 are preferable over the other instruments with less understood measurement models. Future work is required to fully examine the measurement invariance of these instruments in epilepsy and related populations (Meredith, 1993).

Our study had a number of strengths. Not only was it conducted in a large sample of patients, but psychiatric diagnosis was confirmed via gold-standard neuropsychiatric assessment occurring in a multi-disciplinary VEM setting. The convergent validity between the full SCL-90-R and the abbreviated forms was examined using multiple statistical approaches, which included a comparison to clinical diagnosis and symptomatology measured using the HADS. While psychiatric diagnosis was informed by the dominant nosology, a formal semistructured interview was not used. Future research aimed at validating the SCL-90-R and its abbreviated forms is required to confirm predictive validity of the instruments against a formal diagnosis derived from such means. While we investigated the measurement models of the SCL-90-R and its short forms, a more complete analysis of the latent variable structure of these instruments was beyond the scope of this study. Future work might more comprehensively investigate the covariance structure of these instruments, with a particular focus on demonstrating measurement invariance across different VEM diagnostic groups. An important consideration is that our study was limited by the clinical diagnoses observed during VEM. As such, we were limited to validating the SCL-90-R and its short forms against the most prevalent diagnostic groups (depressive disorders, anxiety disorders, psychosis, and obsessive-compulsive disorder). While other diagnoses were made, they either had very low prevalence (e.g., specific phobias) or did not map onto any of the SCL-90-R domains (e.g., eating disorders). A direction for future research is to validate multi-domains instruments like the SCL-90-R using a broader representation of psychiatric disorders.

In summary, these results indicate that several abbreviated forms of the SCL-90-R are appropriate for use in clinical and research settings where more lengthy self-report instruments are contraindicated. In settings where a single measure of global psychopathology is warranted, the 9-item SCL-K-9 provides a valid and reliable solution. In situations where depression and anxiety are of research interest, or where screening for these disorders is desired, the BSI-18 provides the most economical and accurate alternative. These instruments perform at a similar level of accuracy to other brief instruments, including the HADS and NDDI-E. Our results indicate the SCL-90-R and its short forms do not perform well in screening to psychosis or obsessive-compulsive disorder, and we were unable to shed light on how well these instruments screen for other psychiatric diagnoses. In any screening application, it is important to note that, while the SCL-90-R and its abbreviated forms are useful for screening out depression and anxiety, they are not very useful for screening *in* these disorders. Nevertheless, they might be useful for prioritising resources in settings where formal neuropsychiatric assessment is of limited availability.

## Acknowledgements

None

## Data availability

All data and code are available by contacting the corresponding author.

## References

Aben, I., Verhey, F., Lousberg, R., Lodder, J., & Honig, A. (2002). Validity of the beck depression inventory, hospital anxiety and depression scale, SCL-90, and hamilton depression rating scale as screening instruments for depression in stroke patients. Psychosomatics, 43(5), 386–393.

Agrawal, S., Turco, L., Goswami, S., Faulkner, M., & Singh, S. (2015). Yield of monitoring in an adult epilepsy monitoring unit (P2. 097). In: AAN Enterprises.

American Psychiatric Association. (2000). Diagnostic and statistical manual of mental disorders: DSM-IV-TR: American Psychiatric Pub.

American Psychiatric Association. (2013). Diagnostic and statistical manual of mental disorders (DSM-5®): American Psychiatric Pub.

Bentler, P. M. (1990). Comparative fit indexes in structural models. Psychological bulletin, 107(2), 238.

Bentler, P. M. (1995). EQS structural equations program manual (Vol. 6): Multivariate software Encino, CA.

Blumer, D., Montouris, G., & Hermann, B. (1995). Psychiatric morbidity in seizure patients on a neurodiagnostic monitoring unit. The Journal of neuropsychiatry and clinical neurosciences, 7(4), 445–456.

Brophy, C. J., Norvell, N. K., & Kiluk, D. J. (1988). An examination of the factor structure and convergent and discriminant validity of the SCL-90R in an outpatient clinic population. Journal of Personality Assessment, 52(2), 334–340.

Cohen, J. (1988). Statistical power analysis for the behavioral sciences (2nd ed.). United States of America: Lawrence Erlbaum Associates.

Cohen, M. L., Testa, S. M., Pritchard, J. M., Zhu, J., & Hopp, J. L. (2014). Overlap between dissociation and other psychological characteristics in patients with psychogenic nonepileptic seizures. Epilepsy & Behavior, 34, 47–49.

Cyr, J., McKenna-Foley, J., & Peacock, E. (1985). Factor structure of the SCL-90-R: is there one? Journal of Personality Assessment, 49(6), 571–578.

de Oliveira, G. N., Lessa, J. M. K., Gonçalves, A. P., Portela, E. J., Sander, J. W., & Teixeira, A. L. (2014). Screening for depression in people with epilepsy: comparative study among neurological disorders depression inventory for epilepsy (NDDI-E), hospital anxiety and depression scale depression subscale (HADS-D), and Beck depression inventory (BDI). Epilepsy & Behavior, 34, 50–54.

Derogatis, L. (1992). SCL-90-R, administration, scoring and procedures manual-II for the R(evised) version and other instruments of the Psychopathology Rating Scale Series. Townson: Clinical Psychometric Research.

Derogatis, L. (1993). The Brief Symptom Inventory (BSI): administration, scoring, and procedures manual. Minneapolis: National Computer Services.

Derogatis, L. (2000). The Brief-Symptom-Inventory-18 (BSI-18): administration, scoring and procedures manual. Minneapolis: National Computer Services.

Derogatis, L., & Savitz, K. The SCL-90-R and the Brief Symptom Inventory (BSI) in Primary Care. Maruish, ME. Handbook of psychological assessment in primary care settings, 297–334.

Diprose, W., Sundram, F., & Menkes, D. B. (2016). Psychiatric comorbidity in psychogenic nonepileptic seizures compared with epilepsy. Epilepsy & Behavior, 56, 123–130.

Endermann, M. (2005). The Brief Symptom Inventory (BSI) as a screening tool for psychological disorders in patients with epilepsy and mild intellectual disabilities in residential care. Epilepsy & Behavior, 7(1), 85–94.

Fisher, R. S., Acevedo, C., Arzimanoglou, A., Bogacz, A., Cross, J. H., Elger, C. E., † Glynn, M. (2014). ILAE official report: a practical clinical definition of epilepsy. Epilepsia, 55(4), 475–482.

Gilliam, F. G., Barry, J. J., Hermann, B. P., Meador, K. J., Vahle, V., & Kanner, A. M. (2006). Rapid detection of major depression in epilepsy: a multicentre study. The Lancet Neurology, 5(5), 399–405.

Green, S. B., & Yang, Y. (2009). Reliability of summed item scores using structural equation modeling: An alternative to coefficient alpha. Psychometrika, 74(1), 155–167.

Hardt, J., & Gerbershagen, H. U. (2001). Cross-validation of the SCL-27: A short psychometric screening instrument for chronic pain patients. European Journal of Pain, 5(2), 187–197.

Harfst, T., Koch, U., Kurtz von Aschoff, C., Nutzinger, D., Rüddel, H., & Schulz, H. (2002). Entwicklung und Validierung einer Kurzform der Symptom Checklist-90-R. DRV-Schriften, 33, 71–73.

Hopp, J., Anderson, K., Krumholz, A., Gruber-Baldini, A., & Shulman, L. (2012). Psychogenic seizures and psychogenic movement disorders: are they the same patients? Epilepsy & Behavior, 25(4), 666–669.

Jones, J. E., Hermann, B. P., Woodard, J. L., Barry, J. J., Gilliam, F., Kanner, A. M., & Meador, K. J. (2005). Screening for major depression in epilepsy with common selfreport depression inventories. Epilepsia, 46(5), 731–735.

Jöreskog, K. G. (1969). A general approach to confirmatory maximum likelihood factor analysis. Psychometrika, 34(2), 183–202.

Karouni, M., Arulthas, S., Larsson, P. G., Rytter, E., Johannessen, S. I., & Landmark, C. J. (2010). Psychiatric comorbidity in patients with epilepsy: a population-based study. European journal of clinical pharmacology, 66(11), 1151–1160.

Klaghofer, R., & Brähler, E. (2001). Konstruktion und Teststatistische Prüfung einer Kurzform der SCL-90–R. Zeitschrift für Klinische Psychologie, Psychiatrie und Psychotherapie.

Leathem, J. M., & Babbage, D. R. (2000). Affective disorders after traumatic brain injury: cautions in the use of the Symptom Checklist-90-R. The Journal of head trauma rehabilitation, 15(6), 1246–1255.

Lesser, R. P. (1996). Psychogenic seizures. Neurology, 46(6), 1499–1507.

Meredith, W. (1993). Measurement invariance, factor analysis and factorial invariance. Psychometrika, 58(4), 525–543.

Müller, J. M., Postert, C., Beyer, T., Furniss, T., & Achtergarde, S. (2010). Comparison of eleven short versions of the Symptom Checklist 90-Revised (SCL-90-R) for use in the assessment of general psychopathology. Journal of Psychopathology and Behavioral Assessment, 32(2), 246–254.

Pearson, K. (1895). Notes on Regression and Inheritance in the Case of Two Parents. Proceedings of the Royal Society of London, 58, 240–242.

Prinz, U., Nutzinger, D. O., Schulz, H., Petermann, F., Braukhaus, C., & Andreas, S. (2013). Comparative psychometric analyses of the SCL-90-R and its short versions in patients with affective disorders. BMC psychiatry, 13(1), 104.

R Core Team. (2019). R: A language and environment for statistical computing. Vienna, Austria: R Foundation for Statistical Computing.

Rai, D., Kerr, M. P., McManus, S., Jordanova, V., Lewis, G., & Brugha, T. S. (2012). Epilepsy and psychiatric comorbidity: a nationally representative population-based study. Epilepsia, 53(6), 1095–1103.

Rauter, U. K., Leonard, C. E., & Swett, C. P. (1996). SCL-90-R factor structure in an acute, involuntary, adult psychiatric inpatient sample. Journal of Clinical Psychology, 52(6), 625–629.

Rayner, G. (2017). The contribution of cognitive networks to depression in epilepsy. Epilepsy currents, 17(2), 78–83.

Robin, X., Turck, N., Hainard, A., Tiberti, N., Lisacek, F., Sanchez, J.-C., & Müller, M. (2011). pROC: an open-source package for R and S+ to analyze and compare ROC curves. BMC bioinformatics, 12(1), 77.

Rosseel, Y. (2012). Lavaan: An R package for structural equation modeling and more. Version 0.5-12 (BETA). Journal of Statistical Software, 48(2), 1–36.

Ruis, C., van den Berg, E., van Stralen, H. E., Huenges Wajer, I. M., Biessels, G. J., Kappelle, L. J., … van Zandvoort, M. J. (2014). Symptom Checklist 90–Revised in neurological outpatients. Journal of clinical and experimental neuropsychology, 36(2), 170–177.

Schmitz, N., Hartkamp, N., Kiuse, J., Franke, G., Reister, G., & Tress, W. (2000). The symptom check-list-90-R (SCL-90-R): a German validation study. Quality of Life Research, 9(2), 185–193.

Sereda, Y., & Dembitskyi, S. (2016). Validity assessment of the symptom checklist SCL-90-R and shortened versions for the general population in Ukraine. BMC psychiatry, 16(1), 300.

Steiger, J., & Lind, J. (1980). Statistically-based tests for the number of common factors. Paper presented at the Annual Spring Meeting of the Psychometric Society, Iowa City.

Swinkels, W. A. M., Kuyk, J. v., Van Dyck, R., & Spinhoven, P. (2005). Psychiatric comorbidity in epilepsy. Epilepsy & Behavior, 7(1), 37–50.

Tellez-Zenteno, J. F., Patten, S. B., Jetté, N., Williams, J., & Wiebe, S. (2007). Psychiatric comorbidity in epilepsy: a population-based analysis. Epilepsia, 48(12), 2336–2344.

Tucker, L. R., & Lewis, C. (1973). A reliability coefficient for maximum likelihood factor analysis. Psychometrika, 38(1), 1–10.

Urbán, R., Kun, B., Farkas, J., Paksi, B., Kökönyei, G., Unoka, Z., … Demetrovics, Z. (2014) .Bifactor structural model of symptom checklists: SCL-90-R and Brief Symptom Inventory (BSI) in a non-clinical community sample. Psychiatry research, 216(1), 146–154.

Vassend, O., & Skrondal, A. (1999). The problem of structural indeterminacy in multidimensional symptom report instruments. The case of SCL-90-R. Behaviour Research and Therapy, 37(7), 685–701.

Weatherburn, C. J., Heath, C. A., Mercer, S. W., & Guthrie, B. (2017). Physical and mental health comorbidities of epilepsy: population-based cross-sectional analysis of 1.5 million people in Scotland. Seizure, 45, 125–131.

West, S. G., Taylor, A. B., & Wu, W. (2012). Model fit and model selection in structural equation modeling. In R. Hoyle (Ed.), Handbook of structural equation modeling (pp. 209–231). New York: Guilford Press.

Wiebe, S., Rose, K., Derry, P., & McLachlan, R. (1997). Outcome assessment in epilepsy: comparative responsiveness of quality of life and psychosocial instruments. Epilepsia, 38(4), 430–438.

Youden, W. J. (1950). Index for rating diagnostic tests. Cancer, 3(1), 32–35.

Zigmond, A. S., & Snaith, R. P. (1983). The hospital anxiety and depression scale. Acta psychiatrica scandinavica, 67(6), 361–370.

