## Appendix A for "Abbreviated assessment of psychopathology in patients with suspected seizure disorders" (from another version of this preprint: version 1, 2019-06-20)

*Table A1.* Scale reliability coefficients

| *SCL-90-R* | | | *SCL-27* | | | *SCL-14* | | | *SCL-K-9* | | | *BSI-53* | | | *BSI-18* | | |
|---|---|---|---|---|---|---|---|---|---|---|---|---|---|---|---|---|---|
| *Domain* | *Items* | *ω* | *Domain* | *Items* | *ω* | *Domain* | *Items* | *ω* | *Domain* | *Items* | *ω* | *Domain* | *Items* | *ω* | *Domain* | *Items* | *ω* |
| SOM | 12 | .88 | DEP | 4 | .85 | DEP | 6 | .89 | TOT | 9 | .86 | SOM | 7 | .86 | SOM | 6 | .81 |
| OCD | 10 | .91 | DYS | 4 | .84 | PHO | 4 | .75 | | | | OCD | 6 | .88 | DEP | 6 | .91 |
| SEN | 9 | .91 | VEG | 6 | .84 | SOM | 4 | .78 | | | | SEN | 4 | .84 | ANX | 6 | .85 |
| DEP | 13 | .93 | AGO | 5 | .80 | TOT | 14 | .89 | | | | DEP | 6 | .91 | TOT | 18 | .94 |
| ANX | 10 | .91 | SOC | 4 | .85 | | | | | | | ANX | 6 | .87 | | | |
| HOS | 6 | .88 | MIS | 4 | .81 | | | | | | | HOS | 5 | .85 | | | |
| PHO | 7 | .87 | TOT | 27 | .95 | | | | | | | PHO | 5 | .88 | | | |
| PAR | 6 | .84 | | | | | | | | | | PAR | 5 | .81 | | | |
| PSY | 10 | .86 | | | | | | | | | | PSY | 5 | .79 | | | |
| TOT | 90 | .97 | | | | | | | | | | TOT | 53 | .97 | | | |

**Note:** *SOM* = somatisation, *OCD* = obsessive compulsive, *SEN* = interpersonal sensitivity, *DEP* = depression, *ANX* = anxiety, *HOS* = hostility, *PHO* = phobic anxiety, *PAR* = paranoid ideation, *PSY* = psychoticism, *DYS* = dysthymic symptoms, *VEG* = vegetative symptoms, *AGO* = agoraphobia, *SOC* = symptoms of social phobia, *MIS* = symptoms of mistrust, *TOT* = reliability for the total scale, *ω* = McDonald's omega, *items* = number of items in each scale.
